# STING activation counters glioblastoma by vascular alteration and immune surveillance

**DOI:** 10.1101/2023.09.03.556091

**Authors:** Justin V. Joseph, Mathilde S. Blaavand, Huiqiang Cai, Fabienne Vernejoul, Rasmus W. Knopper, Thomas B. Lindhardt, Kristian A. Skipper, Esben Axelgaard, Line Reinert, Jacob G. Mikkelsen, Per Borghammer, Søren E. Degn, Eric Perouzel, Henrik Hager, Brian Hansen, Joanna M. Kalucka, Mikkel Vendelbo, Søren R. Paludan, Martin K. Thomsen

**Affiliations:** Department of Biomedicine, Aarhus University, Aarhus, Denmark; InvivoGen, Toulouse, France; Center of Functionally Integrative Neuroscience, Department of Clinical Medicine, Aarhus University, Aarhus, Denmark; Department of Pathology, Aarhus University Hospital, Aarhus, Denmark; Department of Nuclear Medicine & PET Centre, Aarhus University Hospital, Aarhus, Denmark; Aarhus Institute of Advanced Studies (AIAS), Aarhus University, Aarhus, Denmark

**Keywords:** GBM, mouse models, CRISPR, immunotherapy, STING agonist

## Abstract

Glioblastoma (GBM) is an aggressive brain tumor with a median survival of 15 months and has limited treatment options. Immunotherapy with checkpoint inhibitors has shown minimal efficacy in combating GBM, and large clinical trials have failed. New immunotherapy approaches and a deeper understanding of immune surveillance of GBM are needed to advance treatment options for this devastating disease. In this study, we used two preclinical models of GBM: orthotopically delivering either GBM stem cells or employing CRISPR-mediated tumorigenesis by adeno-associated virus, to establish immunologically proficient and non-inflamed tumors, respectively. After tumor development, the innate immune system was activated through long-term STING activation by a pharmacological agonist, which reduced tumor progression and prolonged survival. Recruitment and activation of cytotoxic T-cells were detected in the tumors, and T-cell specificity towards the cancer cells was observed. Interestingly, prolonged STING activation altered the tumor vasculature, inducing hypoxia and activation of VEGFR, as measured by a kinome array and VEGF expression. Combination treatment with anti-PD1 did not provide a synergistic effect, indicating that STING activation alone is sufficient to activate immune surveillance and hinder tumor development through vascular disruption. These results guide future studies to refine innate immune activation as a treatment approach for GBM, in combination with anti-VEGF to impede tumor progression and induce an immunological response against the tumor.

## 1. Introduction

Glioma is a malignancy in the brain involving the glial cells, and glioblastoma (GBM) is a grade IV glioma with limited treatment options and an extremely poor prognosis associated with a median survival of just under 15 months ^1^. Gliomas are divided into different subtypes, with isocitrate dehydrogenase (IDH)-mutated gliomas being classified as low risk ^2^. Most GBMs arise in elderly patients and are IDH-intact. Molecular classification of GBM based on transcriptomic analysis has emerged as a new tool complementing the traditional pathology-based description ^3, 4^. These approaches have identified three major molecular subtypes of GBM namely-classical, proneural and mesenchymal tumors ^5^. Proneural tumors often harbor mutations in PDGFR and TP53, whereas mesenchymal GBMs often contain mutations in NF1 and PTEN ^4, 5^. In general, the standard of care for GBM involves surgical intervention followed by radiation and chemotherapy with temozolomide ^1^. Despite extensive research and investment into the advancement of glioma treatment, little progress has been made ^6^. Therefore, there is an urgent need to explore new treatment possibilities to prolong survival and eventually cure this devastating disease.

In recent years, immunotherapy has been explored for the treatment of various cancer types and has greatly improved treatment of certain cancers, especially melanoma and lung carcinoma ^7–9^. Similar approaches has been tested in GBM and checkpoint inhibitors such as anti-PD1/PD-L1 or CTLA4 have shown remarkable results in preclinical models of GBM ^10–14^. However, these promising results had no translational relevance as it failed to show efficacy in glioma patients ^15–17^. Combination treatment with conventional irradiation or chemotherapy together with checkpoint inhibitors has also resulted in disappointing results ^18^. The lack of efficacy could partly be attributed to the heterogeneity of the tumors and the reduced drug delivery over the blood-brain barrier (BBB), even though the BBB is often disrupted in the tumor milieu ^19^. Furthermore, gliomas are often classified as cold tumors with minimal infiltration of cytotoxic T-cells and few genetic mutations to present neoantigens for immunological recognition ^20^. Efforts are being made to modulate the tumor milieu to attract cytotoxic T-cells, but also to generate further mutations in the cancer cells to increase the possibility of immunological recognition ^20^.

Immune activation has shown promising results in preclinical models of cancer. Towards this direction the DNA-sensing receptor cyclic GMP–AMP synthase (cGAS) and its downstream signaling effector stimulator of interferon genes (STING) have been explored, despite belonging to the innate immunity pathway for recognition of DNA in the cytosol from viral infections ^21^. Activation of the STING pathway induces the expression of type 1-IFN and a plethora of interferon-stimulated genes (ISG), including cytokines with chemotactic effects on T-cells, such as CXCL10 ^21, 22^. The binding of DNA to cGAS generates the signal molecule cGAMP, which binds to STING and activates the pathway. The effects of cGAMP or analogs have been tested in preclinical models of cancer and have shown reduced tumor growth and tumor clearance ^23^. Combination treatment with checkpoint inhibitors has shown even more promising results, and clinical trials are ongoing with these combination approaches ^24, 25^. STING agonists have preferably been administered intratumorally to achieve a high local concentration, and this has also been tested in an orthotopic model of GBM resulting in a reduced tumor load ^26^. Another approach has been systemic delivery with low doses over extended period to ensure immune activation, as intra-tumoral delivery is not always feasible ^27^.

In this study, we systemically administered a cGAMP analog, cAIMP bisphosphorothioate and difluorinated (CL656) ^28^, which is currently in clinical trial (NCT04592484) and has increased stability in vivo. To evaluate the effect on GBM, we orthotopically implanted the GBM stem cell line 005 into the striatum of mice and treated established tumors for three weeks. STING activation reduced tumor progression and increased survival by increasing T-cell activation. Similar results were obtained in CRISPR-induced GBM mouse models, even though these tumors harbored fewer cytotoxic T-cells. It was observed that prolonged activation of STING also altered blood vessels in the tumors, resulting in larger but fewer vessels leading to increased hemorrhages and hypoxia in the tumor milieu. Overall, this study shows that STING activation increases tumor surveillance partly through immune activation and through alteration of vasculature. Together these results pave the way to future studies to refine combination treatments with anti-VEGF and checkpoint inhibitors.

## 2. Methods

### 2.1. Cell work

The cell line 005 which was a kind gift from prof Samuel D. Rabkin (Harvard medical school) and is described in ^29^. These cells were propagated as free-floating spheroids in neural stem cell medium (NSM) which is composed of-Neurobasal A-Medium (Gibco) supplemented with 2% B27 supplement (Gibco), 20ng/ml EGF (peprotech), 20 ng/ml bFGF (peprotech), 1% pen/strep and 1% L-glutamine (Gibco Life Technologies) and heparin. Sub culturing was done once in 3-4 days and involved pelleting and washing cells with PBS followed by incubation with accutase (Sigma-Aldrich) and repeated pipetting in NSM medium to dissociate the spheroids. The single cell suspension was seeded into a flask with fresh NSM in appropriate culture ratio. Cells were maintained at 37°C in a humidified atmosphere with 5% CO2.

### 2.2. Animals

The mice used in the experiments were bred and housed at the animal facility of Aarhus University. All animal experiments were conducted in accordance with the protocol approved by the Danish Animal Experiments Inspectorate (license no. 2018-15-0201-01589). The animals were kept in pathogen-free facilities following institutional guidelines. Wildtype C57bl/6j mice were purchased from Taconic, while B6J.129(B6N)-Gt(ROSA)26Sortm1(CAG-cas9*,-EGFP)Fezh/J and STINGgt/gt (STING-Goldenticket; C57BL/6J-Tmem173gt/J) mice were obtained from Jackson Laboratories. The mice were anesthetized at 8-10 weeks of age by intraperitoneal injection of anesthetics containing medetomidine hydrochloride (1 mg/mL), midazolam (5 mg/mL), and butorphanol (10 mg/mL). Cells and viral delivery of the sgRNA/Cre transgene cassette were achieved by orthotopic injections into the striatum of the mouse brain. Using stereotaxic surgery, a total of 3 µl of cells (20000 cells in total) or 2 µl of AAV (5*10^9vg) were injected at the coordinate’s A/P +1, M/L -2, D/V −3.5 over 10 minutes. The mice recovered on a heating pad and received atipamezole (5 mg/kg) as an antidote and Carprofen (0.05 mg/10g) as an analgesic.

### 2.3. Drug delivery

cAIMP bisphosphorothioate and difluorinated (cAIM(PS)2 Difluor(Rp/Sp) (CL656)) were injected intraperitoneally in saline three times a week for three weeks. Two doses of CL656 were tested, 0.2 µg/g or 0.5 µg/g, and thereafter, 0.2 µg/g was used for all experiments (InvivoGen, tlrl-nacairs-2). Anti-PD1 (InvivoGen, pdl1-mab15-1) was injected intraperitoneally at 100 ng/g twice a week for two weeks. Pimonidazole was administered intraperitoneally at 60 µg/g in saline 1 hour prior to termination (Hypoxyprobe).

### 2.4. PET/MR imaging

Animals underwent anatomical magnetic resonance imaging (MRI) to follow disease progression, and a subset received 18[F]-Fluoroethyl-L-tyrosine (FET) for positron emission tomography (PET). Mice were anesthetized with isoflurane, which was maintained throughout the scanning procedure using a respiration mask. A bolus of FET (approximately 15 MBq/animal) was administered via tail vein injection. PET scanning was performed 60 minutes after the injection, followed by MRI scanning.

### 2.5. Ex-vivo Magnetic Resonance Imaging

Mice were perfusion-fixed with 4% PFA following which the brain was harvested intact and stored in 4% PFA for 3-4 weeks. Before imaging, samples were washed in PBS for 24 hours and securely placed in an MRI-suitable tube filled with fluorinert (FC-770, 3M inc.) as described previously ^30^. High resolution MRI was performed using a Bruker Biospec 9.4T preclinical MRI system (Bruker Biospin, Ettlingen, Germany). High resolution B0-mapping was performed to aid field shimming, performed with Bruker’s MapShim. The structural data were acquired using a RARE sequence at an in-plane resolution of 50 µm × 50 µm and a slice thickness of 200 µm (80 slices in total). For diffusion kurtosis imaging (DKI) data was acquired at 200 µm isotropic resolution using a segmented (8 segments) diffusion weighted echo planar imaging sequence. Five unweighted images were acquired in each slice for normalization along with 30 isotropically distributed encoding directions at each of 4 non-zero b-values (0.5, 1.0, 1.8, 2.5 ms/µm2). Diffusion timings were δ/Δ = 7/16 ms, and remaining parameters were 10 avs, TE=39.2 ms, TR = 4000 ms, BW= 277 kHz, 80 axial slices. DKI analysis has been described previously ^31, 32^. Briefly, data was Rician noise floor adjusted, denoised and corrected for Gibbs-ringing before fitting to the DKI signal equation (Matlab, The Mathworks, USA). Mean diffusivity (MD) and mean kurtosis (MK) were calculated for each sample and regions of interest (ROIs) were manually drawn. In this manner, the ipsilateral and contralateral striatum was outlined in each sample and used for ROI analysis. All sample preparation, data acquisition, analysis, and delineation were performed blinded.

### 2.6. AAV production

Specific single guide RNAs (sgRNAs) (Table 1S) were cloned into an AAV2 backbone. Two vectors were generated, both containing sgRNAs for Pten and Trp53 under S7K and U6 promoter, respectively. Furthermore, sgRNAs for either Nf1 or Rb1 were included in the vector under the H1 promoter. The expression of Cre was driven by the GFAP promoter to restrict expression to astrocytes. AAV particles were generated using a helper plasmid and the PHP.eB serotype (Addgene). Virus production was conducted in HEK293T cells ^33^ and purified as described previously ^34^. Virus titer was determined by viral genome (vg) quantification using qPCR as previously described ^33^, and 5*10^9 vg was injected per animal to induce glioma formation.

### 2.7. Histology and Immunohistochemistry

Histological sections were cut from PFA-fixed tissue embedded in paraffin. Antigen retrieval was carried out at 100°C in citrate buffer with a pH of 6 for 20 minutes. Endogenous peroxidase was blocked by incubation with 0.3% hydrogen peroxide for 10 minutes. Sections were then blocked in 5% BSA in TBS containing 0.1% Tween 20. The following primary antibodies were used: F4/80 (AB6640), CD4 (CS25229), CD8 (CS98941), CD105 (RD, AF1320), Viperin (Millipore, MABF106), PD1 (CST, 84651), and Anti-pimonidazole (Hpi, MAb1). Counterstaining was performed using hematoxylin. H&E staining was performed using standard protocols.

### 2.8. Quantification of CD105 staining

For morphometric analyses, 5 random optical fields (10X magnification) per tumor section were taken by a Zeiss Axioplan upright microscope (Carl Zeiss, Munich, Germany) and analyzed using Fiji (Image J) as described previously ^35^. The measurements of vessel number were performed by counting number of vessels segments within the tumor area, and tumor vessel area was measured as total CD105+ area, and expressed as a percentage of the total tumor area.

### 2.9. DNA, RNA and Protein isolation from animal tissue

Total DNA and RNA were isolated from the samples using the Qiagen AllPrep DNA/RNA kit according to the manufacturer’s protocol. Protein lysates were prepared using M-PER mammalian extraction buffer (Pierce, Thermo Scientific) supplemented with 1× protease and phosphatase inhibitor cocktail. Samples underwent sonication to homogenize and fragment genomic DNA.

### Sanger sequencing and indel analysis

DNA isolated from the CRISPR-induced tumors was analyzed for mutations in the target genes. The regions of interest were amplified by PCR using DreamTaq Green PCR Master Mix (ThermoFisher) according to the manufacturer’s recommendations (see Table S1 for primer sequences). Gel-purified PCR products were subjected to Sanger sequencing, and data were analyzed using the ICE webtool (Synthego) to identify indel formations.

### 2.10. Real-time quantitative PCR

20 ng of total RNA isolated from tumors was used for gene expression analysis. Gene-specific primers were used (Table S1) together with Brilliant III Ultra-Fast SYBR Green QPCR Master Mix (Agilent Technologies), following the manufacturer’s recommendations. Data were analyzed using the ΔΔCT method, and expression of Actin was used to normalize gene expression levels.

### 2.11. Western blotting

Isolated protein extracts were subjected to SDS-PAGE immunoblotting, and membranes were blocked with either 2.5% BSA (Sigma-Aldrich) or 5% dry milk (BD) in TBS containing 0.1% Tween20 (Sigma-Aldrich). The following primary antibodies were used: pSrc (CST, 2101), pLyn (CST, 2731), pStat1 (CST, 7649), Sting (CST, 13647), cleaved caspase 3 (CST, 9661), Isg15 (CST, 2743), and Vinculin (Sigma, V9131). Anti-rabbit or anti-mouse horseradish peroxidase-conjugated secondary antibodies were used for development (Jackson ImmunoResearch).

### 2.12. Kinome analysis

The PamGene chips were used according to the manufacturer’s protocol. Four individual protein samples isolated from GBM 4 hours after final PBS or CL656 treatment were processed for assay analysis according to the manufacturer’s instructions. Both tyrosine kinase assay (PTK) and serine-threonine kinase (STK) arrays were performed. Signal intensities were analysed in the BioNavigator software (PamGene) and expressed as log2 fold changes versus PBS control. The differential kinases among the groups were analysed by the Upstream Kinase Analysis (UKA) tool, and the kinome tree was made by the CORAL tool (htp://phanstiel-lab.med.unc.edu/CORAL/).

### 2.13. Statistical analyses

A Log-rank (Mantel-Cox) test was used for statistical analysis of the Kaplan-Meier survival curve. An unpaired t-test was used for analysis of two groups by using GraphPad Prism. A p-value < 0.05 was considered statistically significant between the two groups.

## 3. Results

### 3.1. STING agonist activates an innate immune response in the CNS

STING agonist has been administered in various mouse models and clinical trials are undergoing ^24^. The stability of STING agonist in vivo is variable and we first sought to address the activation of STING pathway in the CNS by the cAIMP bisphosphorothioate and difluorinated (cAIM^(PS)^2 Difluor(Rp/Sp) (CL656)) agonist. Mice were treated 3 times a week for 3 weeks with 0.2 or 0.5 mg/kg and the weight was monitored. Mice receiving 0.2mg/kg did not diverge in their weight from PBS control group, whereas mice receiving the high dose lost weight in the last week (Fig S1). To evaluate the activation of STING pathway in the CNS, mice were treated with 0.2mg/kg CL656 for nine times over 3 weeks and samples were taken 4 hours after the last treatment. Histological examination showed no difference between PBS or CL656 treatment, except expression of the interferon stimulated gene Viperin. Here staining was found throughout the brain tissues but with increased expression on the endothelial cells (Fig 1a). Analysis of Viperin expression in the human protein atlas shows similar expression in malignant human CNS samples (Fig S2). ELISA for Ifnβ confirmed elevated levels in serum 4 hours after treatment and the expression of interferon stimulated gene (ISGs) in the CNS were increased (Fig 1b, c). This was confirmed by Western blotting (WB) that showed increased levels of phosphorylated Stat1 and Viperin in the treated groups (Fig 1d). Interestingly, the apoptotic marker cleaved caspase 3 was detected in samples receiving CL656 (Fig 1d), indicating a cytotoxic effect of STING activation in the CNS. Overall, systemic delivery of the STING agonist CL656 activates the STING pathway in CNS, with specific upregulation of ISGs but also with limited cytotoxic effect.

**Figure 1:**
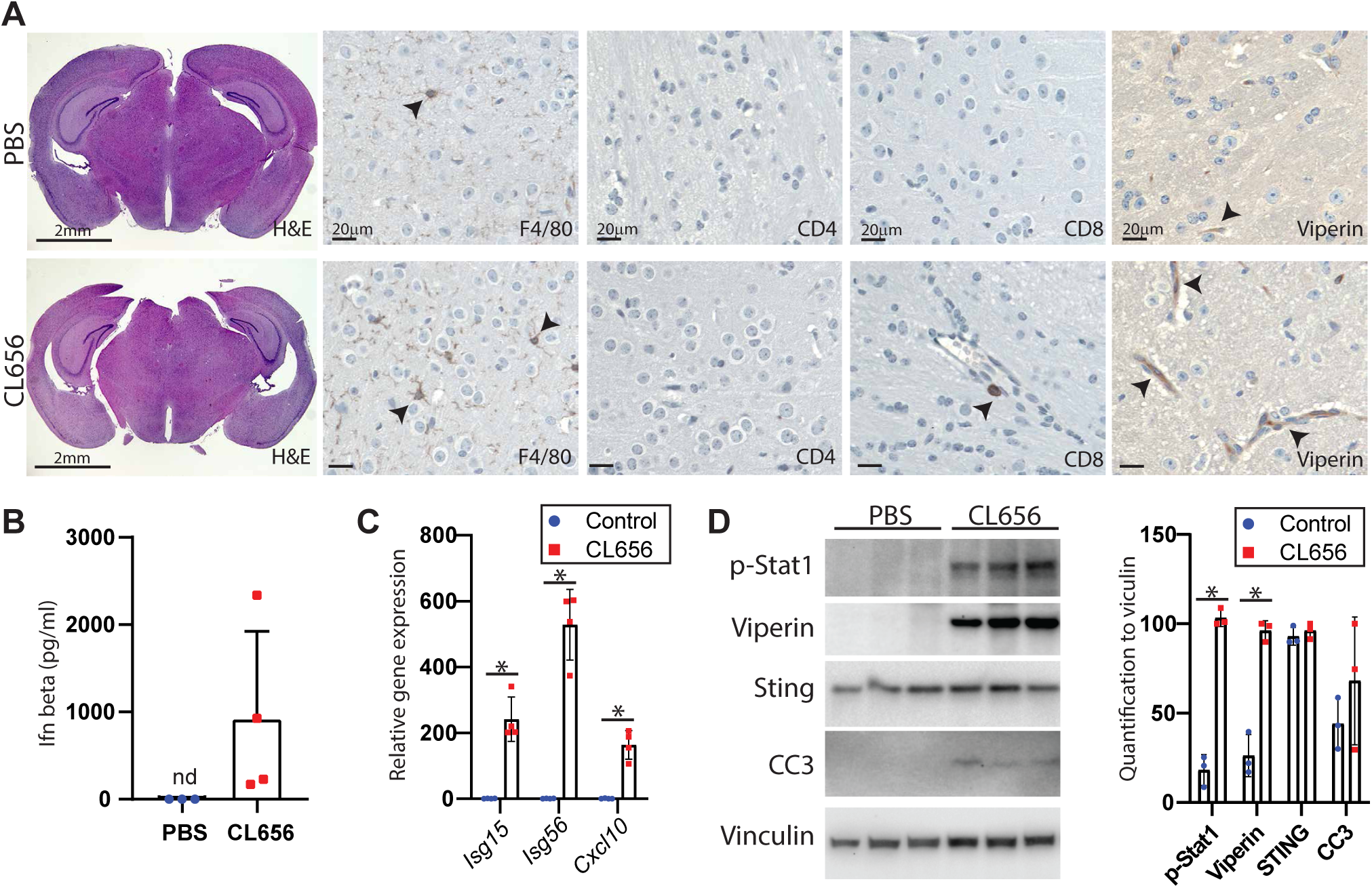
ISG expression in the brain after systemic STING activation. Mice received nine doses of STING agonist (CL656) or PBS during a 3-week period. After the last dose, mice were sacrificed 4 hours post-treatment, and samples were collected for analysis. A) IHC was performed on brain sections of PBS-or CL656-treated mice (n=4). B) Ifn-beta was measured in the serum of the mice (n=4, nd=not-detected). C) Total RNA was isolated from brain tissues, and gene expression of Isg15, Isg56, and Cxcl10 was measured from CL656-or PBS-treated mice (n=4, *=p<0.05). D) Western blot was performed on protein isolated from the brain of treated mice. Band intensity was measured and normalized to Vinculin (n=3, *=p<0.05).

### 3.2. Long-term activation of STING pathway hampers tumor development

Immune therapy towards glioma has showed contradictory results ^15, 17^. To assess the effect of modified STING agonist CL656 in GBM we generated an orthotopic GBM model by implanting the mouse GBM stem cell line 005 into the striatum of the mouse. MRI scan was performed three weeks post implantation to confirm tumor formation. Hereafter, mice received nine doses of CL656 over a period of three weeks followed by a second in vivo MRI scan at the completion of treatment to evaluate treatment response on tumor growth (Fig 2a, b). Assessment of MRI scans revealed decreased tumor volume and diameter in the CL656 treated group compared to PBS treated mice (Fig 2c). Application of a PET tracer, 8-fluoride-fluoro-ethyl-tyrosine (FET) confirmed the reduced tumor size after STING activation (Fig S3). These findings were confirmed by macroscopic examination of the tumors after the last treatment, as the 005 cells expressed GFP (Fig 2d). However, at humane endpoint the tumors had achieved similar size, but the pace of tumor progression was delayed in mice that received CL656 treatment, which resulted in a prolonged survival (Fig 2e-f, S4). Together, CL656 treatment over three weeks hampered tumor progression and prolonged survival.

**Figure 2:**
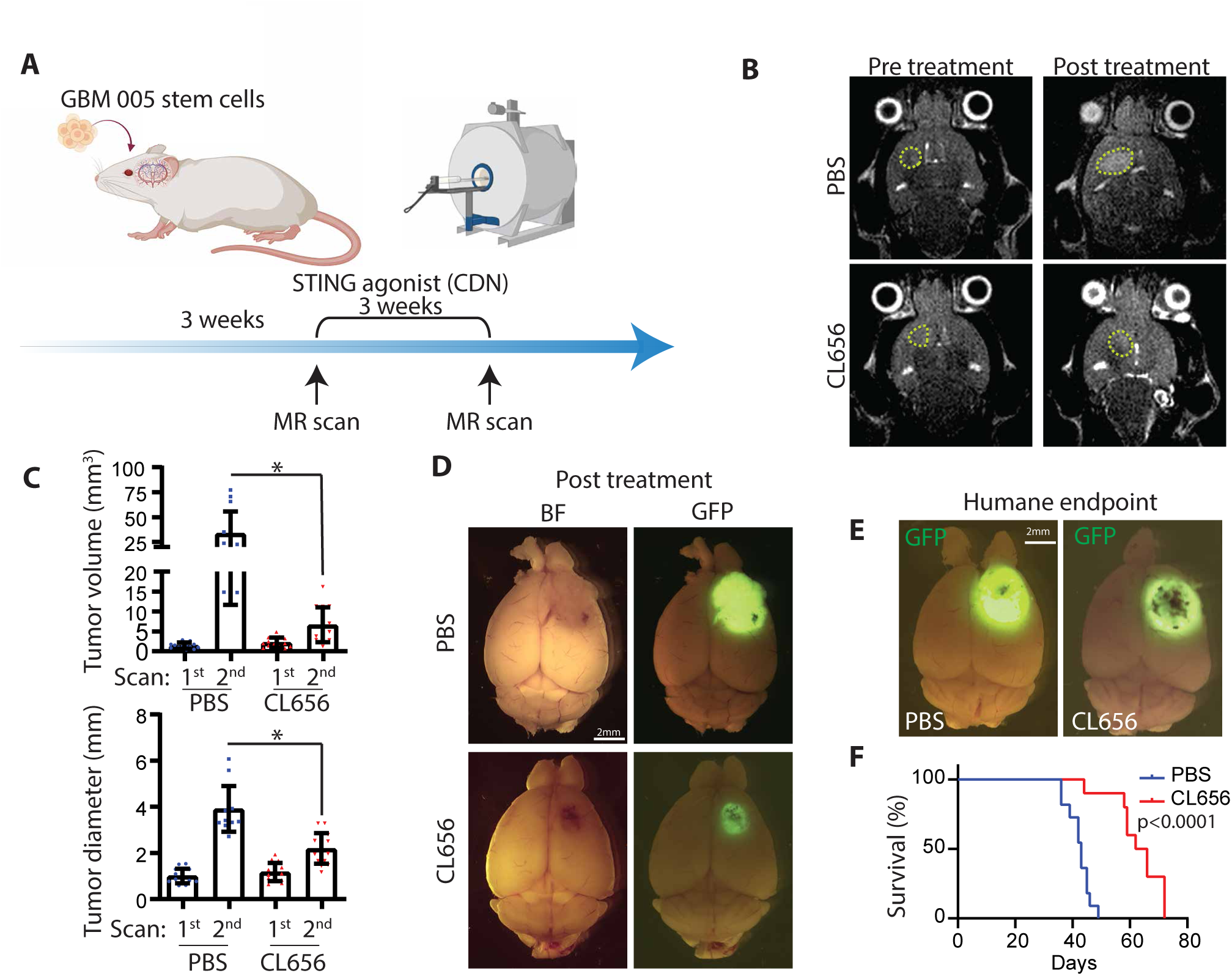
STING activation increased survival in a pre-clinical model of GBM. A) Mice were orthotopically implanted with the murine GBM cell line 005 into the striatum. Three weeks later, an MRI scan was performed to confirm tumor development. Subsequently, mice were treated with STING agonist (CL656) nine times over 3 weeks, followed by an MRI scan to monitor tumor response. B) MRI scans before and after treatment with PBS or CL656. A representative image is shown, and yellow dash line mark the tumor (n>10). C) Tumor diameter and volume were calculated by MRI scans between the groups (n>8-10, *=p<0.05, Pre-treatment scan indicated as 1^st^ and Post-treatment scan is indicated as 2^nd^). D) A subgroup of the mice was killed 4 hours after the last treatment. Tumors from PBS-or CL656-treated mice were assessed by bright field (BF) and fluorescence microscopy for GFP (n>5, representative images are shown). E) Mice were followed until the humane endpoint, and fluorescence microscopy was used to assess the tumors between the two groups (n>10, representative images are shown). F) The survival of PBS-or CL656-treated mice was assessed at the humane endpoint (n=10).

### 3.3. STING agonist activates T-cells and mediates tumor cytotoxicity

The antitumor property of STING agonist is often driven by activation of cytotoxic T-cells, but it has also been reported to have a direct effect on the cancer cells. To assess the direct effect of STING agonist on 005 tumor cells, these cells were treated with CL656 which led to an induction of Isg15 expression due to STING activation (Fig S5a). Next, 005 cells were implanted into the striatum of STING deficient mice followed by treatment with CL656 or PBS. Mice were scanned before and after treatment, but no difference was observed between the two groups of mice indicating tumor-extrinsic STING dependency of CL656 to induce anti-tumor effect (Fig S5b, c). Furthermore, the tumor growth pattern and survival of the mice in both the control and treatment groups was similar revealing that CL656 had little or no direct effect on the tumor cells (Fig S5d, e).

This result steered us to address the effect of CL656 on non-tumor cells with focus on cytotoxic T-cells. To assess the presence of CD8 positive T-cells in the tumor, histological sections were evaluated after final treatment. Staining for CD8 showed that cytotoxic T-cells were only confined to the tumor tissues and not distributed in the normal brain parenchyma (Fig 3a). Staining for PD1 showed positive cells in the tumors marking activated T-cells (Fig 3b). Interestingly, quantification of CD8 or PD1 positive cells in the tumor showed similar scores between CL656 or PBS treated mice (Fig 3c). However, similar analyses were done at humane endpoint and here CL656 treated mice showed elevated levels of PD1 positive cells but similar amount of CD8 cells (Fig 3e-f). To assess T-cell tumor surveillance, a group of mice received anti-PD1 or CL656 in combination with anti-PD1 (Fig 3g). Anti-PD1 treatment prolonged survival but combination treatment with CL656 had minor effect (Fig 3h). This was reflected in increased amount of CD8 and PD1 positive T-cells in the tumor of anti-PD1 treated mice. The combination treatment increased the amount of PD1 positive cells as was observed for CL656 treatment alone (Fig3 e, f, i). Collectively, these data indicate that CL656 activates cytotoxic T-cells in the GBM.

**Figure 3:**
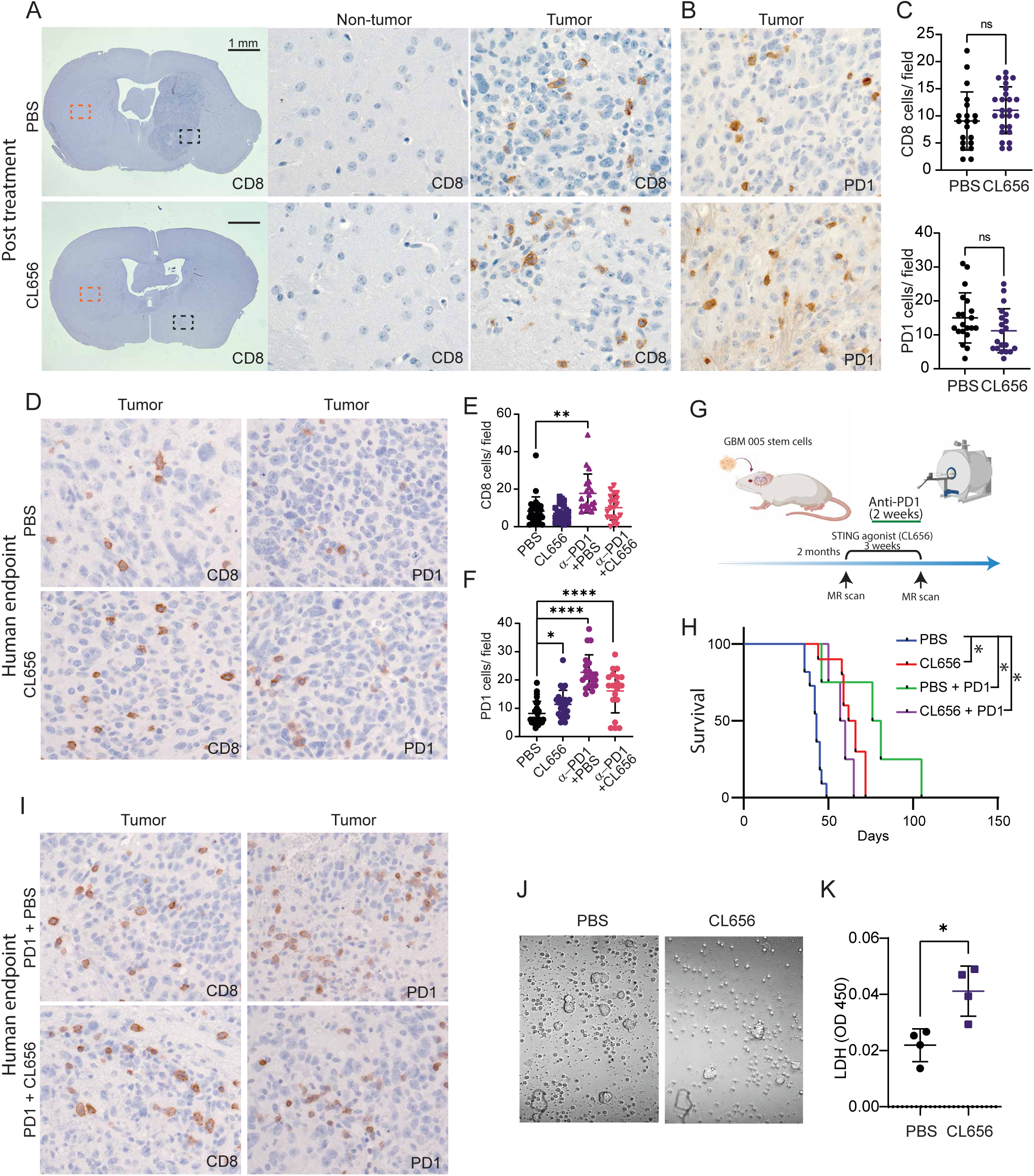
STING Agonist Enhances T-cell Activation and Eradication of Tumor Cells. A) IHC for CD8 was performed on tissue sections from the brain 4 hours after the last treatment with PBS or CL656 (n=5). The red box marks the non-tumor area, and the black box marks the tumor. B) IHC for activated T-cells by PD1 expression was assessed in tumor tissues (n=5). C) Quantification of CD8 and PD1-positive cells on tumor samples derived from PBS or CL655 treated animals (n=25 fields, ns= not significant). D) IHC for CD8 and PD1 on tumor sections at humane endpoint (n=10, a representative image is shown). E, F) Quantification of positive CD8 or PD1-positive cells was assessed on tumor sections at humane endpoint (n=25 fields, *=p<0.05, **=p<0.01, ****=p<0.0001). G) A subset of mice received four doses of anti-PD1 over two weeks, in combination with either PBS or CL656. H) The survival of mice treated with anti-PD1 in combination with CL656 or PBS was compared to mice treated with anti-PD1 alone (n=10 for mono therapy, n=4 for combination treatment). I) Immunohistochemistry (IHC) was performed for CD8 and PD1 on tumor sections from mice that received anti-PD1 in combination with either PBS or CL656 (n=4; a representative image is shown). J) CD8-positive T-cells were isolated from tumor-bearing mice that had been treated with either PBS or CL656. The isolated T-cells were co-cultured with 005 tumor cells in vitro for 6 hours, and bright-field images were taken (30:1 (T-cell/Tumor cells)). K) The killing of tumor cells by the isolated CD8-positive T-cells was assessed by the LDH assay from the co-culture experiment after 6 hours (n=4; * indicates p<0.05).

To assess the cytotoxic effect of T-cells towards 005 tumor cells, mice were inoculated with 005 cells and treated with CL656 or PBS as previously. After the last treatment, CD8 positive T-cells were isolated and enriched (Fig S6). These CD8 T-cells were co-cultured with 005 tumor cells in vitro. Cell death was assessed and here CD8 positive cells isolated from mice treated with CL656 showed increased capability to kill 005 cells when compared with T-cells from PBS treated mice (Fig3j, k). This data shows that CL656 induced activation of STING increases the activity of cytotoxic T-cells towards the glioma cells.

### 3.4. Vascular alteration by STING activation

To investigate further the possible anti-cancer effect of STING agonist on GBM progression, we performed a kinome array on protein extracted from tumor samples after final treatment to assess activity of nearly 340 kinases. This analysis revealed that a number of cancer related kinases had reduced activity after CL656 treatment (Fig 4a-d, S7). This included Src and Lyn kinases, which both work upstream of Myc and we confirmed downregulation of these two kinases by Western blotting in CL656 treated tumors (Fig 4b, c). Interestingly, several of the kinases were upregulated including the FLT family hereunder FLT1, which codes for Vegfr1 (Fig 4d). The ligand for Vegfr1 is Vegfa, which promotes angiogenesis and qPCR data showed increase levels of Vegfa and partial upregulation of the hypoxia induced gene Eno1 in CL656 treated samples (Fig 4e). The upregulation of Vegf pathway coincides with increased hemorrhage observed in CL656 treated mice, which were visualized by MRI scans (Fig S8a) and macroscopic tumor examinations (Fig 4f). Histological analysis revealed hemorrhage within the tumor and staining for CD105 as an endothelial marker showed altered vasculature between the two groups of mice (Fig 4g). Quantification of vascular area showed an increase in the CL656 treated group, but the vessel number was decreased, presumably because of increased vessel diameter (Fig 4h). However, this alteration of the vessels was only observed in the tumor and not in the normal tissues (Fig S8b). Overall, these results collectively shows that STING agonist impairs the tumor milieu by alterations of blood vessels and activation of the Vegf pathway.

**Figure 4:**
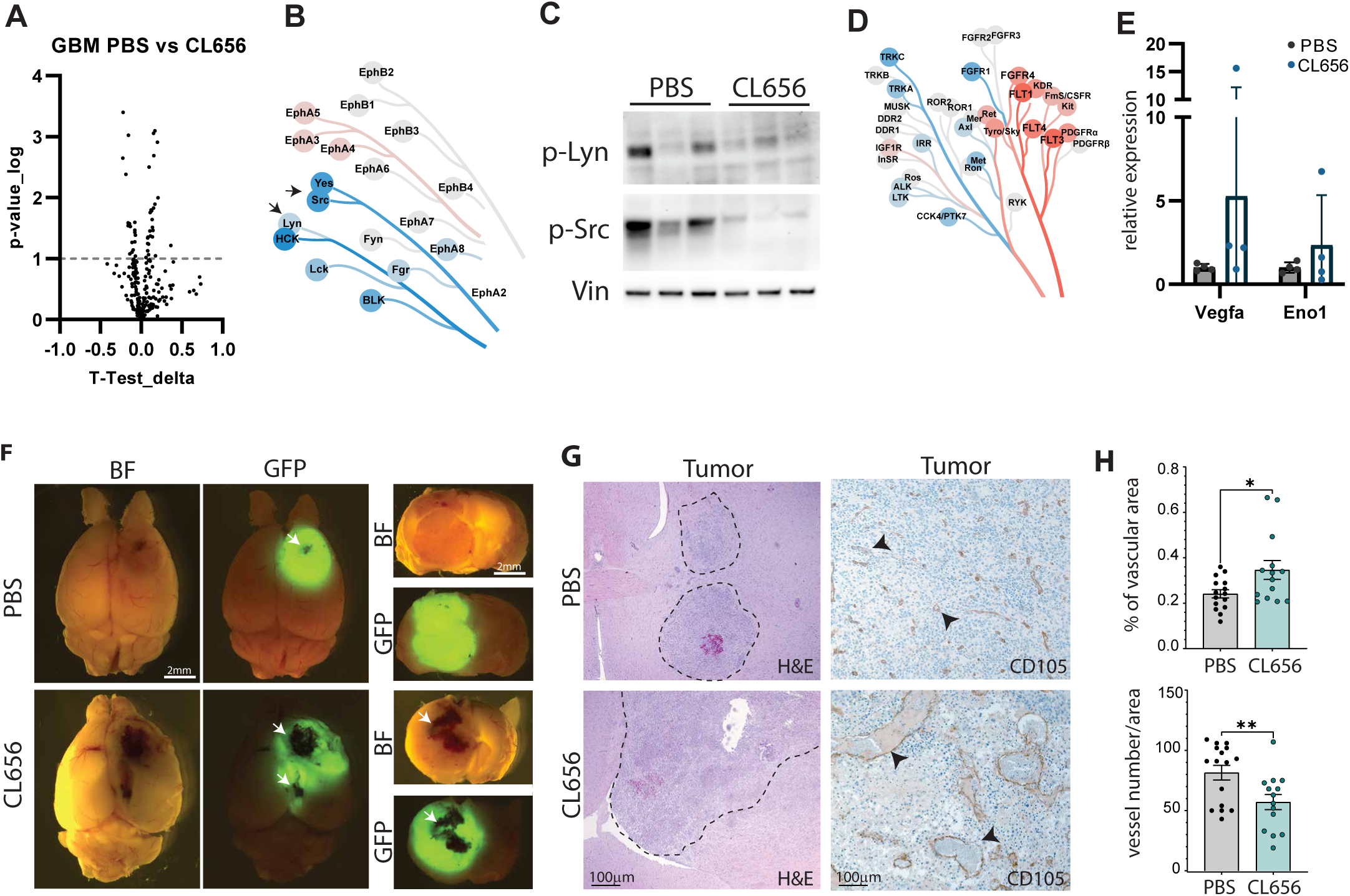
STING agonist alters tumor vasculature. A) A kinome array for 340 kinases was performed on protein isolated 4 hours after the last treatment of PBS or CL656 (n=4). The phosphorylation of target peptides between the two groups is displayed. B) A coral tree (median final score) between PBS and CL656 treatment displays a decrease in Src Family Kinases. C) Western blot for phosphorylated Lyn and Src on protein lysate from tumors 4 hours after the last treatment of PBS or CL656 (n=3). D) A coral tree (median final score) between PBS and CL656 treatment displays an increase in the Tyrosine Kinase Receptor Flt/VEGFR Family. E) Gene expression of Vegfa and Eno1 in tumor tissues 4 hours after the last treatment of PBS or CL656 (n=4). F) Tumors from PBS or CL656-treated mice were assessed by bright field and fluorescence microscopy at humane endpoint. White arrowheads mark large hemorrhages (n>5; representative images are shown). G) Tissue sections of tumors at humane endpoint were stained with H&E or the endothelial cell marker CD105. The dashed line marks the tumor border, and black arrowheads mark blood vessels positive for CD105 (n=5; representative images are shown). H) Quantification of CD105 staining was performed to evaluate vascular area and vessel number in tumor tissues at humane endpoint between PBS-and CL656-treated mice (n=25 fields/group, * = p < 0.05, ** = p < 0.01).

### 3.5. Administration of STING agonist on CRISPR generated model of glioblastoma

Cell line derived xenograft models of cancer tend to be immunogenic and suitable for immune therapy, whereas spontaneously occurring tumors are less immunogenic ^36^. To evaluate CL656 implication on glioma originating in the healthy organ, CRISPR was applied to induce cancer promoting mutations in a subset of astrocytes. Two models of glioma were established with mutation in either Rb1 or Nf1 in combination with loss of Trp53 and Pten. The two models had different latency, where Rb1 tumors could be detected by MRI 8 months after induction, Nf1 tumors were visible 10 weeks after induction (Fig 5a, S9, S10a, b). Mice receiving AAV with Cre expression did not develop tumors (Fig S10c, d). When tumor was detected by MRI mice underwent 9 treatments over 3 weeks with CL656 or PBS as control. The CL656 treatment did not alter tumor size in mice containing Rb1 mutations over the three weeks of treatment, probably due to the slow progression of the tumor (Fig S9). However, when the whole brain was scanned after the last treatment by ex-vivo diffusion kurtosis imaging (DKI), analysis revealed altered microstructure in the tumors from mice that had received CL656 treatment (Fig S9d, e).

**Figure 5:**
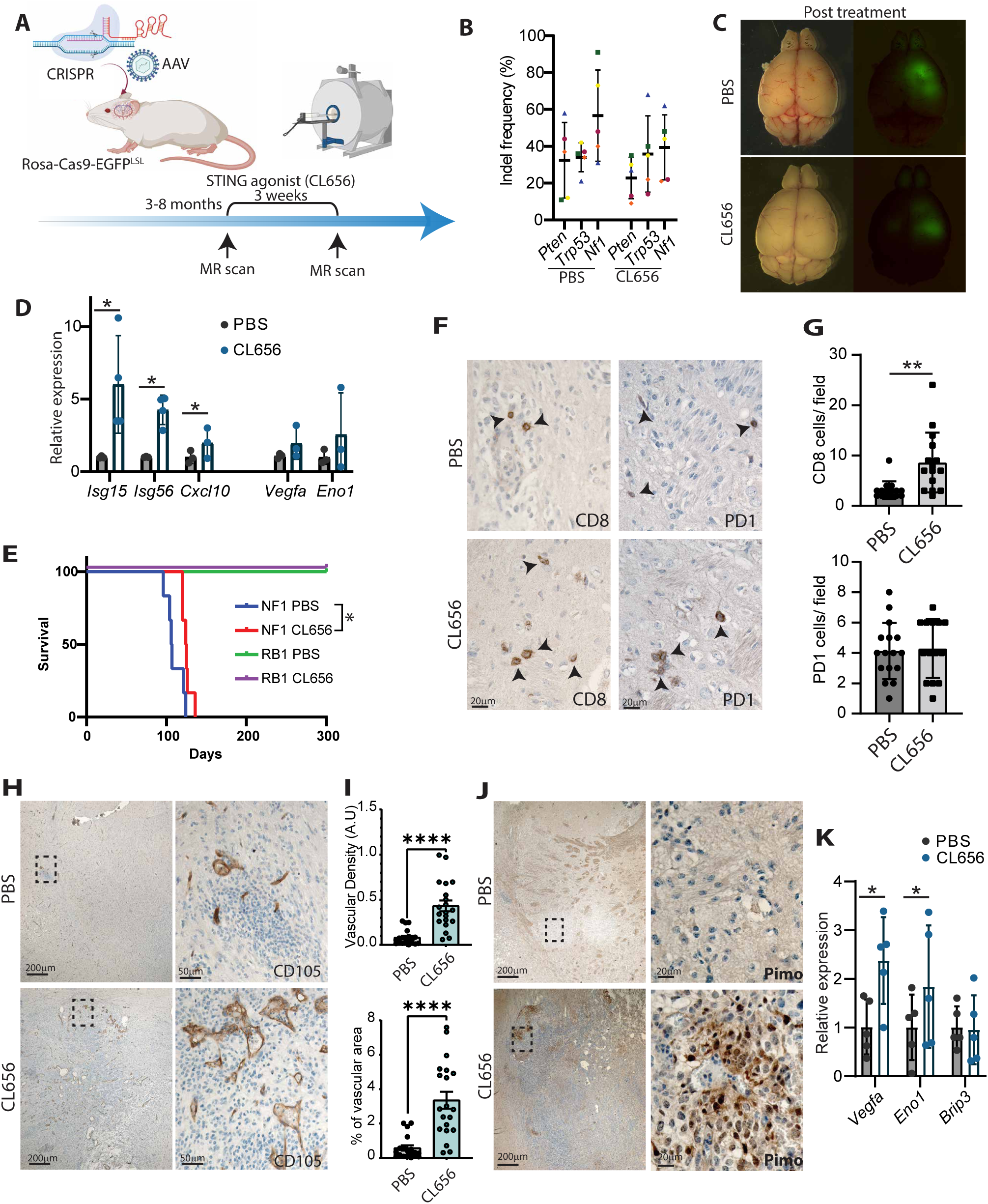
Increased T-cells infiltration and hypoxia in a CRISPR model of GBM upon STING activation. A) GBM was induced by AAV delivery of CRISPR guides to a Cas9 transgene mouse. sgRNAs targeting Nf1 or Rb1 in combination with sgRNAs for Pten and Trp53 were applied. Tumor progression was monitored by MRI scans, and when the tumors reached approximately 1 mm in diameter, treatment with PBS or CL656 was conducted nine times over a three-week period. Post-treatment, an MRI scan was conducted to evaluate treatment response. B) CRISPR-induced mutations were assessed in tumor samples from mice treated with PBS or CL656. InDel frequency for Nf1, Trp53, and Pten was analyzed by ICE-webtool (n=5). C) Tumors from PBS or CL656-treated mice were assessed four hours after the last treatment by bright-field and fluorescence microscopy (n>5, representative images are shown). D) Quantitative expression of type 1-IFN-induced, and hypoxia-related genes was analyzed four hours post-last treatment (n=3, * = p < 0.05). E) The survival of mice bearing GBM induced by CRISPR was followed post treatment with either PBS or CL656 (n=8, *p<0.05). F) Histological sections of Nf1 tumors, taken 4 hours after the last treatment, were stained for CD8 or PD1. Black arrowheads indicate positive cells (n>5). G) Quantification of CD8 and PD1 positive cells in Nf1 deficient tumors from mice treated with PBS or CL656 (n=25 fields, **p<0.01). H) IHC for CD105 was performed on sections from Nf1 deficient tumors at humane endpoint. The dash box indicates the magnified image on the left (n=5, representative image is shown). I) Quantification of CD105 staining was used to evaluate vascular area and vessel density in tumor tissues at humane endpoint between PBS and CL656 treated mice (n=25 fields/group, ****p<0.0001). J) IHC for hypoxia was performed on sections from Nf1 deficient tumors at humane endpoint. One hour prior to termination, mice received pimonidazole. The dash box indicates the magnified image on the left (n=6, representative image is shown). K) Quantitative expression of hypoxia and autophagy-related genes was assessed at humane endpoint (n=5, *p<0.05).

Next, we focused on GBM with Nf1 mutations (Fig 5b), as this group showed a much more rapid tumor progression. A subset of mice was terminated after the last treatment with CL656 (Fig 5c) and expression of interferon stimulated genes was measured on these tumor samples. Increased levels were detected comparable to mice that had been implanted with 005 glioma cells (Fig 5d). Expression of Vegfa and the Hif1 alpha induced gene, Eno1, were also assessed and both genes showed elevated expression levels (Fig 5d). Another group of mice were followed up to humane endpoint and mice that had received CL656 treatment had a slightly prolonged survival (Fig 5e). To evaluate the prolonged survival after CL656 treatment, histological examination was performed to assess the presence of CD8 T-cells and activation by PD1 expression (Fig 5f). Quantification revealed increased amount of CD8 positive cells in the tumors of CL656 treated mice but no difference was found for cells positive for PD1 (Fig 5g). A subset of mice received combination treatment with anti-PD1 and CL656, but no synergetic effect was seen on tumor progression or survival (Fig S11).

As was observed with 005 tumors treated with CL656, the CRIPSR induced tumors had increased presentation of hemorrhage (Fig S10). Histological evaluation revealed changes in vascular structure of CL656 treated mice by CD105 staining (Fig 5h). Quantification confirmed that CL656 treatment caused remarkable increase in vascular density and area (Fig 5i). The functionality of these vessels is unknown, but we assessed the mice for possible hypoxia in the tumors by pimonidazole analysis. Here, mice that had received CL656 showed areas in the tumor that was positive for pimonidazole, revealing that administration of CL656 could induce hypoxia. Similar, increased expression of Vegfa and Eno1 were detected in the tumors of mice that had received STING agonist. Overall, these results reveal that STING agonist have dual implication on GBM by both increasing cytotoxic T-cells and by alteration of the blood vessels within the tumor.

## 4. Discussion

In this study, we have demonstrated that prolonged and systemic pharmacological activation of the STING pathway hampers GBM progression and increases overall survival in mouse models of GBM. Immune therapy targeting GBM has shown limited efficacy in clinical trials, despite remarkable results in pre-clinical models ^15, 17^. Pre-clinical models are often based on synergistic cell lines that are highly immunogenic. In our study, we used the GBM stem cell line 005, which harbors multiple genetic alterations and has been passaged in vitro, resulting in increased mutational burden ^29^. The 005-cell line has previously been shown to be suitable for immune therapy approaches, including checkpoint inhibitors and oncolytic viruses ^11^. We observed efficacy of the STING agonist towards GBM derived from orthotopically implanted 005 cells. To assess the effectiveness of STING agonist in a “cold” tumor, we generated GBM through CRISPR-mediated mutations of Nf1, Trp53, and Pten in astrocytes. This resulted in an aggressive tumor that was immunologically cold with remarkably fewer infiltrating cytotoxic T-cells. Interestingly, treatment of these mice with STING agonist decreased tumor progression and increased survival. Activation of STING in these immunologically cold tumors, resulted in an increased number of cytotoxic T-cells in the tumors, revealing that systemic STING activation alters the glioma milieu towards increased infiltration of lymphocytes.

Activation of STING can be achieved pharmacologically using STING agonists or by inhibiting ENPP1, which degrades cGAMP, or through epigenetic activation of STING ^37–39^. STING agonists can be delivered systemically or intra-tumorally, where the later can achieve high local doses ^39^. This can result in the death of tumor cells and increase the presentation of tumor material. However, activation of STING can also result in the death of bystander cells, such as T-cells ^40^. In this study, we administered systemic STING agonist at a low dose and observed ISG expression in the CNS, but also sporadic cell death. Evaluation of tumor samples harvested immediately post completion of the treatment did not show changes in the number of cytotoxic T-cells, although we have previously observed a decrease in CD8-positive cells in a model of HCC after long-term treatment with STING agonist ^27^. Instead, when GBMs were assessed at humane endpoint, both increased amount of T-cell and activation were observed, indicating an expansion of specific T-cell clones targeting the cancer cells, as seen in human patients with GBM ^41^. Furthermore, it has been reported that endothelial cells activated by STING agonist facilitate increased invasion of T-cells to the underlying tissues ^42^. We also observed Viperin-stained endothelial cells as a marker of type 1-IFN stimulation, suggesting activated endothelial cells in the CNS leading to a possible increase in T-cell trafficking and IFN production ^26, 43^.

Despite the increased amounts of cytotoxic T-cells in gliomas due to STING activation, we did not observe an additive effect of combination treatment with anti-PD1. The combination treatment was given in the last two weeks of treatment to overcome immune suppression as a feedback mechanism to STING activation, as PD-L1 is an interferon-stimulated gene ^44, 45^. Both single treatments with the STING agonist and anti-PD1 showed efficiency, but a potential synergistic effect could not be shown. However, other studies with the 005 GBM stem cell line have revealed prolonged survival by combination and even clearance of tumors ^11^. We speculated that prolonged treatment of anti-PD1 after administration of the STING agonist could have improved efficiency, as we have observed increased tumor surveillance in a mouse model of HCC months after STING stimulation, indicating a T-cell driven response ^27^.

Combination treatment for GBM has been investigated in both preclinical models and clinical trials ^6^. An example is the combination of temozolomide with immunotherapy, here the treatments contradict as temozolomide also negatively affects T-cells ^46^. Therefore, this study focused on anti-PD1 to overcome the feedback mechanism towards STING activation. Others have combined irradiation with immune therapy to increase the presentation of tumor antigens and induction of neoantigens ^20^. In this study we tested two different models of glioma, one with the cell implantation model as an immunogenic tumor and the CRISPR-induced model as a “cold” tumor with few mutations. Interestingly, both models showed an increase in the amount of T-cell recruitment by STING activation and also by anti-PD1 treatment. This shows that immune surveillance is ongoing in the different models of glioma. Despite this, the overall survival is increased only by a few weeks. This steers future studies to focus on additional combination treatments, as immune activation only slows tumor progression, and further intervention is needed to overcome resistance to immunotherapy.

DMXAA was the first STING agonist described and was discovered before STING was identified and was believed to be a vascular targeting agent ^47^. Here we showed that both models of GBM showed alteration in the blood vessel structure in the tumors after treatment with STING agonist. An increase in vessel size was observed, but also a decrease in the number of vessels, indicating that STING activation decreases the number of vessels but enlarges their size. A study by Lee et al. demonstrated that STING activation of endothelial cells has been linked to increased vessel structure and normalization in a model of colon cancer ^48^. What makes our study different is the long-term treatment in a different cancer type compared to the short term intra-tumoral treatment done by Lee et al. Furthermore, GBM is particularly interesting because of its location in the brain parenchyma, insulated by the blood-brain-barrier, and our observation of the modulation of vascularization involving the blood-brain-barrier is thus unique. In a previous study, we performed long-term treatment in a model of HCC but did not observe vascular alterations indicating a tumor type specificity to the observed pattern in vasculature ^27^. The functionality of the altered blood vessels in glioma is unknown, but mice that had been treated with STING agonist displayed increased hemorrhage, hypoxia, and increased expression of hypoxia-related genes. Hypoxia is associated with a worse prognosis in GBM patients, partly through immune suppression and upregulation of VEGF ^49^. Therefore, anti-VEGF therapy has shown efficiency in clinical trials both as monotherapy and in combination with temozolomide ^50, 51^. It is possible that a combination therapy of anti-VEGF and STING agonist could provide beneficial results, as the formation of new blood vessels would be hampered in the tumor milieu.

Immunotherapy has shown great efficacy in subtypes of tumors with high immunological presentation and many immune cells within the tumor ^20^. Activation of STING by different agonists can induce an immunological response and potentially modify the tumor environment towards an immunological state^39^. GBM is a non-immunogenic tumor, but we have shown in this study that prolonged systemic activation of the STING pathway can increase the presence of cytotoxic T-cells in the tumor, resulting in improved survival rates. However, the effects of this treatment are modest, and combining it with anti-PD-1 therapy did not result in an additive effect in this setting. Therefore, future research will focus on combination treatment of STING agonists and anti-VEGF towards GBM to discover new opportunities for the treatment of this devastating cancer.

## Acknowledgments

We would like to thank Savithri Rangarajan for assistance with analysis of Pamgene data, Mette Simonsen for assistance with PET/MRI scanning, Heidi Schou Knudsen for her technical help and Karina Lassen Holm for taking care of our mice. This research was funded by Danish Cancer Society (R231-A13828, R204-A12490), and Ministry of health (4-1612-236/7), Dagmar Marshall Fond, P.A. Messerschmidt og Hustrus Fond, Helge Peetz og Verner og hustru Vilma Peetz legat, Raimond og Dagmar Ringgård-Bohns Fond, The Aarhus University Research Foundation, and Thora og Viggo Grove’s Mindelegat (all to MKT). Danish Cancer Society (R311-A18039) to SRP.

