## Supplementary material for "STING activation counters glioblastoma by vascular alteration and immune surveillance": Sup figures

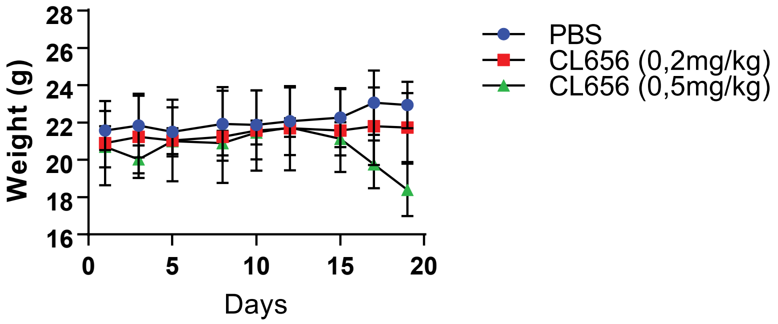

Weight (g)
28
26
24
22
20
18
16
0
5
10
15
20
Days
PBS
CL656 (0,2mg/kg)
CL656 (0,5mg/kg)


Fig S1

**Figure S1: Titration of the STING agonist CL656.**

Mice at 8 weeks of age received nine doses of CL656 at 0.2mg or 0.5mg/kg over three weeks. Control mice were treated with PBS. The weight of the mice was monitored throughout the treatment period (n=6).

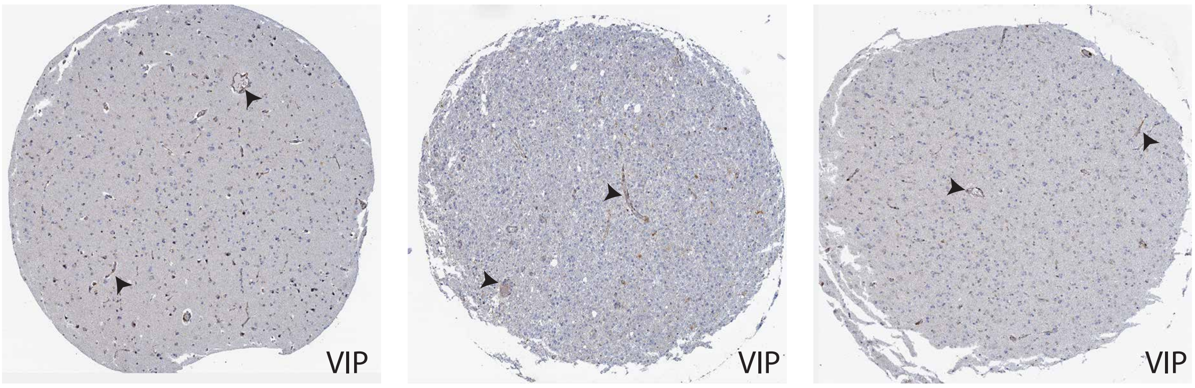


Figure S2

**Figure S2: Expression of Viperin in human glioma**

IHC on human glioma samples shows expression in the endothelial cells. Black arrowheads mark positive blood vessels. Data from the human protein atlas.

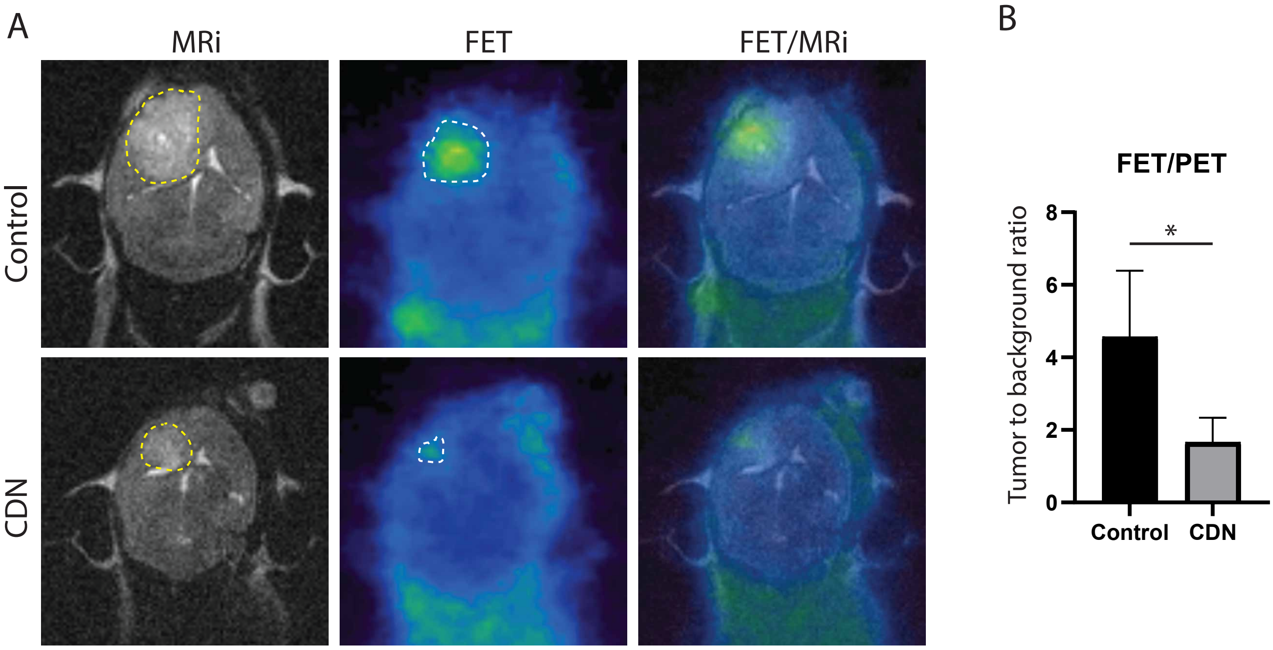
A
MRi
FET
FET/MRi
Control
CDN
B
FET/PET
Tumor to background ratio
8
6
4
2
0
*
Control
CDN

Fig S3

**Figure S3: FET-PET/MRI confirms decreased tumor volume after STING activation.**

A) Mice were orthotopically implanted with the murine GBM cell line 005 into the striatum. Six weeks later, a PET tracer, 8-fluoride-fluoro-ethyl-tyrosine (FET), was used in combination with an MRI scan to visualize tumor lesions in mice treated with PBS or CL656 (n>3).

B) Tumor volumes were calculated based on the FET signal normalized to background (n=3, *=p<0.05).

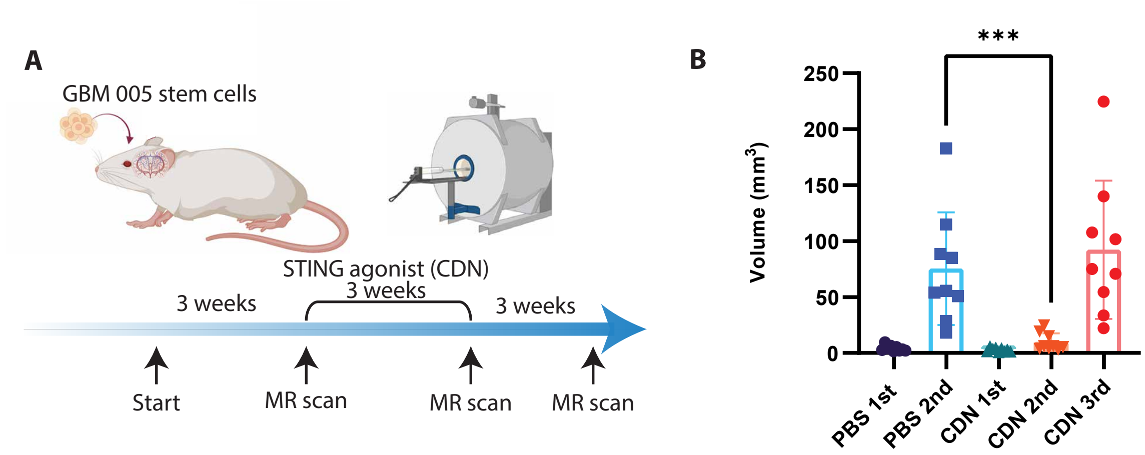
A
GBM 005 stem cells
STING agonist (CDN)
3 weeks
3 weeks
3 weeks
Start
MR scan
MR scan
MR scan
B
***
Volume (mm³)
250
200
150
100
50
0
PBS 1st
PBS 2nd
CDN 1st
CDN 2nd
CDN 3rd

Fig S4

**Figure S4: Delay tumor progression by STING agonist**

A) Mice were orthotopically implanted with the murine GBM cell line 005 into the striatum. Three weeks later, an MRI scan was performed to confirm tumor development. Subsequently, mice were treated with STING agonist (CL656) nine times over 3 weeks, followed by an MRI scan to monitor tumor response. 3 weeks post treatment, mice that had not meet humane endpoint (CL656 treated mice) were scanned for a tried time (n=8).

B) Tumor volumes were calculated by MRI scans between the groups ($n \geq 8$-10, ***=$p<0.001$).

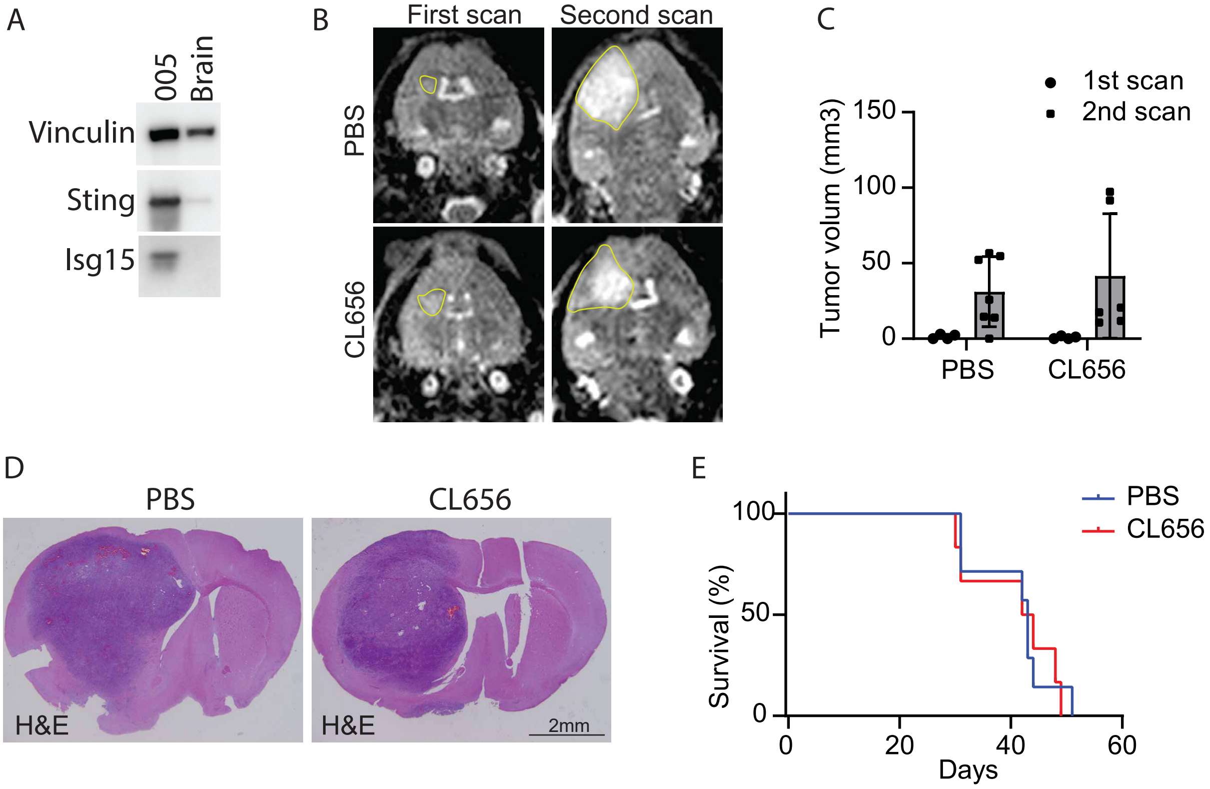
A
005
Brain
Vinculin
Sting
Isg15
B
First scan
Second scan
PBS
CL656
C
1st scan
2nd scan
Tumor volum (mm3)
150
100
50
0
PBS
CL656
D
PBS
CL656
H&E
H&E
2mm
E
PBS
CL656
Survival (%)
100
50
0
0
20
40
60
Days

Fig S5

**Figure S5: Tumor Cell Activation of STING is Dispensable for Tumor Progression**

A) Protein was isolated from the GBM cell line 005 four hours after activation with CL656 and compared with protein lysate from tumor-bearing mice. Western blot was performed for Isg15 and Sting, with Vinculin used as a loading control.

B) STING-deficient mice were orthotopically inoculated with the mouse GBM cell line 005 into the striatum. Three weeks later, an MRI scan was conducted to confirm tumor formation. Subsequently, mice were treated with STING agonist (CL656) nine times over three weeks, followed by an MRI scan to assess tumor response. Yellow lines mark the tumors (n=8).

C) Tumor volume was determined by MRI scans between the groups (n=8).

D) Mice were followed until the humane endpoint, and H&E staining was performed on sections from the tumors (n=8). Representative images are shown.

E) The survival of PBS- or CL656-treated mice was assessed at the humane endpoint (n=8).

A

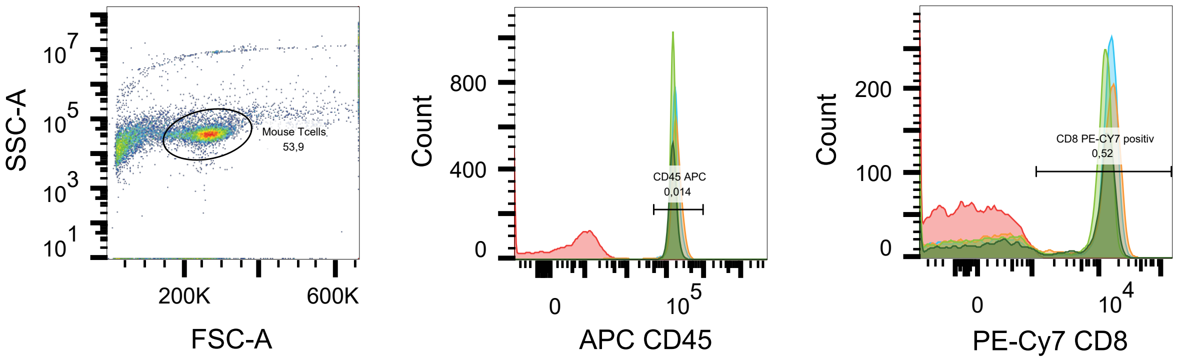


B

| | Sample Name | Subset Name | Count | CD45 APC :: Freq. of Parent |
|---|---|---|---|---|
| (dark green) | Tceller 220421_A5_NEM C.fcs | Mouse Tcells | 5044 | 97,0 |
| (light green) | Tceller 220421_A4_IR C.fcs | Mouse Tcells | 7248 | 97,9 |
| (orange) | Tceller 220421_A3_NEM T.fcs | Mouse Tcells | 7575 | 95,4 |
| (blue) | Tceller 220421_A2_IR T.fcs | Mouse Tcells | 7398 | 96,2 |
| (red) | Tceller 220421_A7_Blank.fcs | Mouse Tcells | 6958 | 0,014 |

C

| | Sample Name | Subset Name | Count | CD8 PE-CY7 positiv :: Freq. of Parent |
|---|---|---|---|---|
| (dark green) | Tceller 220421_A5_NEM C.fcs | Mouse Tcells | 5044 | 62,2 |
| (light green) | Tceller 220421_A4_IR C.fcs | Mouse Tcells | 7248 | 62,7 |
| (orange) | Tceller 220421_A3_NEM T.fcs | Mouse Tcells | 7575 | 62,5 |
| (blue) | Tceller 220421_A2_IR T.fcs | Mouse Tcells | 7398 | 67,1 |
| (red) | Tceller 220421_A7_Blank.fcs | Mouse Tcells | 6958 | 0,52 |

Figure S6

**Figure S6: Isolation of T-cells from tumor-bearing mice.**

Mice were orthotopically inoculated with the mouse GBM cell line 005 into the striatum. Three weeks later, mice were treated with STING agonist (CL656) nine times over 3 weeks. T-cells were isolated from the spleen.

A) T-cells were isolated using beads, and a sample was analyzed by flow cytometry for the expression of CD45 and CD8.

B) Percentage of CD45-positive cells in the samples.

C) Percentage of CD8-positive cells in the samples.

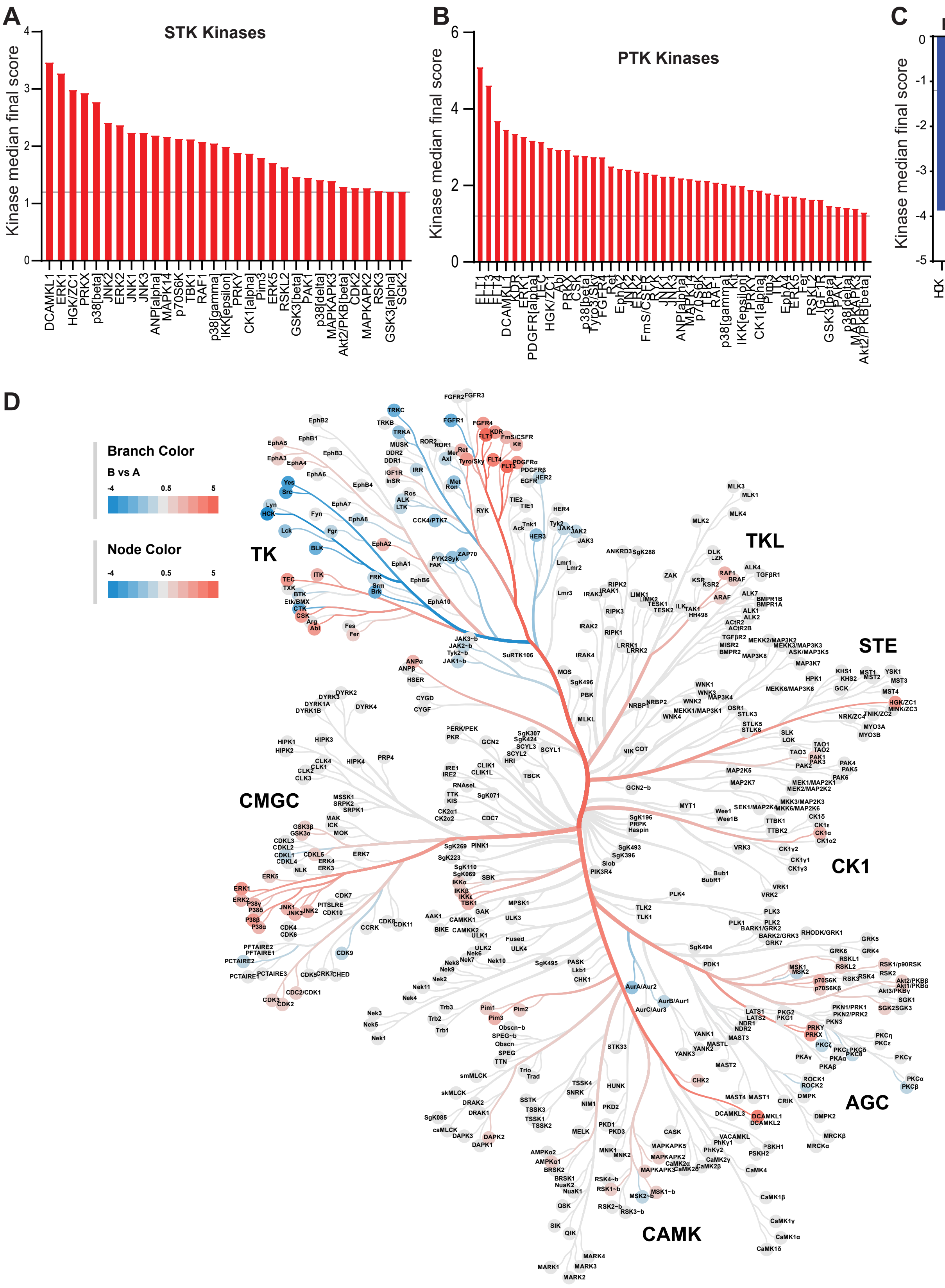
A
STK Kinases
Kinase median final score
DCAMKL1
ERK1
HGK/ZC1
PRKX
p38[beta]
JNK2
ERK2
JNK1
JNK3
ANP[alpha]
MAPK14
p70S6K
TBK1
RAF1
p38[gamma]
IKK[epsilon]
PRKY
CK1[alpha]
Pim3
ERK5
RSKL2
GSK3[beta]
PAK1
p38[delta]
MAPKAPK3
Akt2/PKB[beta]
CDK2
MAPKAPK2
RSK3
GSK3[alpha]
SGK2
B
PTK Kinases
Kinase median final score
C
PTK Kinases
Kinase median final score
HCK
Src
Yes
TRKC
BLK
Brk
CTK
FGFR1
D
Branch Color
B vs A
Node Color
TK
TKL
STE
CMGC
CK1
AGC
CAMK

Fig S7

**Figure S7: Impaired kinase activity by CL656 stimulation.**

A-C) A kinome assay for 340 kinases was performed on protein isolated 4 hours after the last treatment of PBS or CL656 (n=4). The median final score for kinase activity was generated, and significant changes in kinases are shown.

D) A coral tree (median final score) between PBS and CL656 treatment displays all regulated kinases.

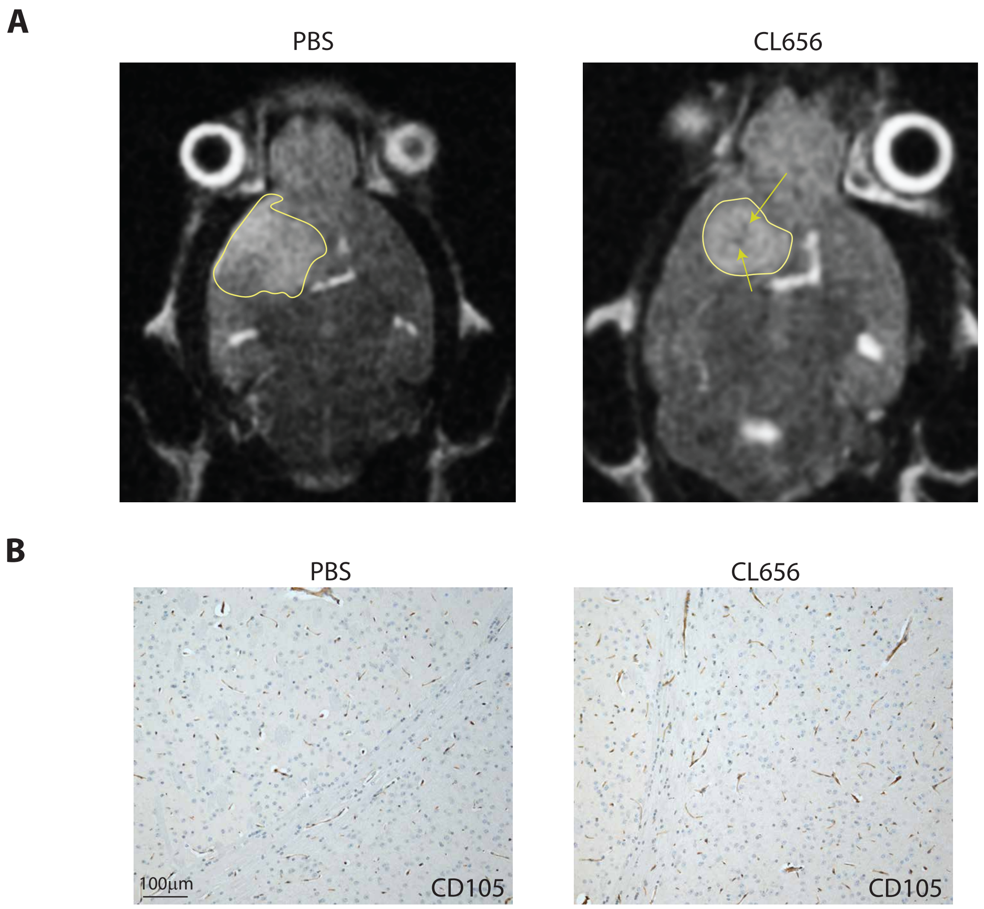


Fig S8

**Figure S8: Hemorrhage Formation after CL656 Treatment**

Mice were orthotopically inoculated with the murine GBM cell line 005 into the striatum. Three weeks later, mice were treated with STING agonist (CL656) nine times over three weeks.

A) MRI scan post-final treatment shows hemorrhage formation in tumors from mice treated with CL656. The yellow line marks the tumors, and the arrows indicate hemorrhage formation.

B) IHC for CD105 on non-tumor tissues from mice treated with PBS or CL656 (n>5).

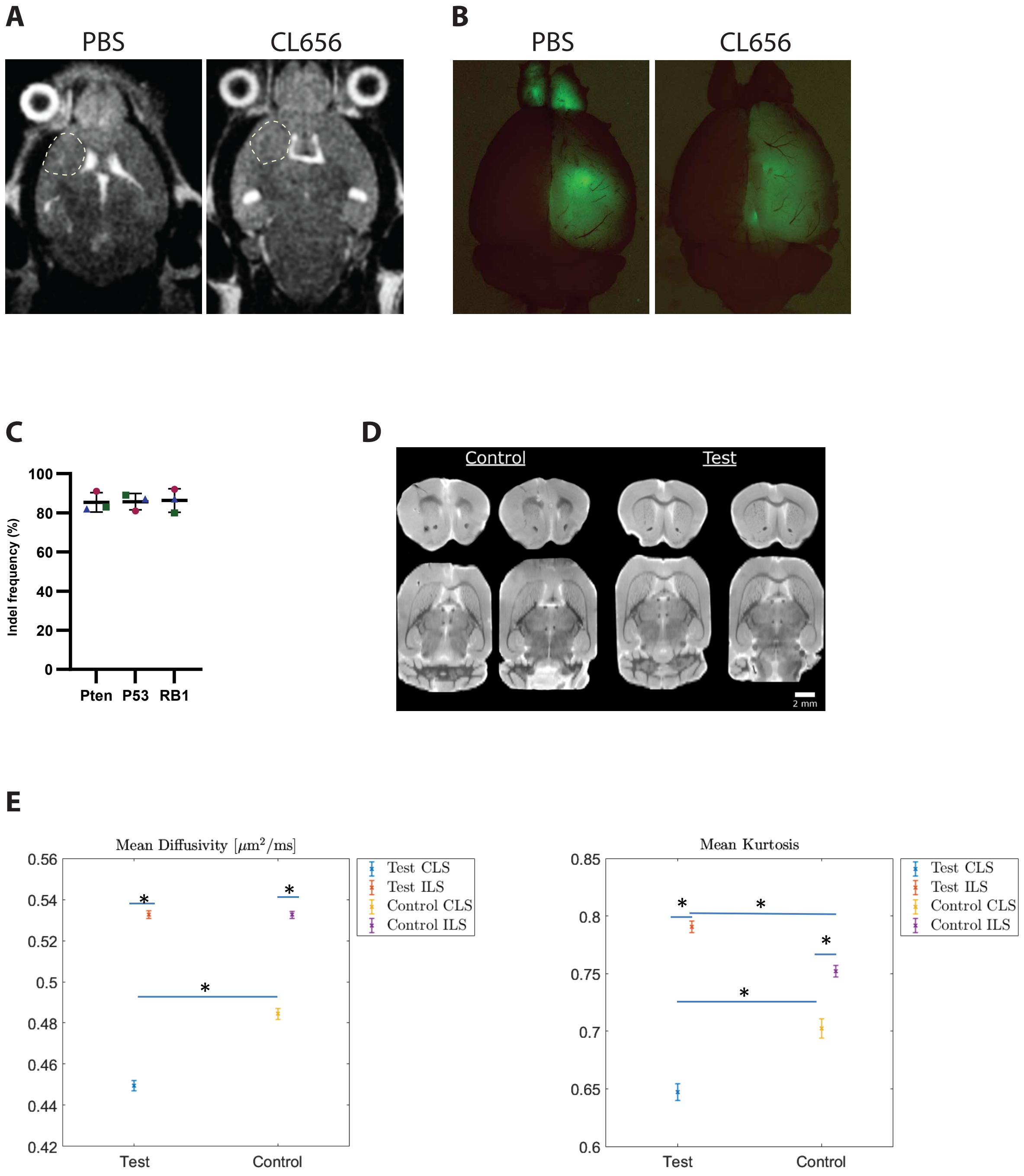
A
PBS
CL656
B
PBS
CL656
C
Indel frequency (%)
100
80
60
40
20
0
Pten
P53
RB1
D
Control
Test
2 mm
E
Mean Diffusivity [μm²/ms]
0.56
0.54
0.52
0.5
0.48
0.46
0.44
0.42
Test
Control
Test CLS
Test ILS
Control CLS
Control ILS
Mean Kurtosis
0.85
0.8
0.75
0.7
0.65
0.6
Test
Control
Test CLS
Test ILS
Control CLS
Control ILS

Fig S9

**Figure S9: Altered tissue architecture after CL656 treatment.**

A) GBM tumors with mutations in Rb1, Pten, and Trp53 were scanned post-treatment. A dashed yellow line marks the tumors (n=10).

B) Rb1 deficient tumors from PBS or CL656-treated mice were assessed by fluorescence microscopy (n>5, representative images are shown).

C) Indel analysis of the tumor tissues for the target genes.

D, E) A subset of whole brains from mice with deficient Rb1 gliomas was isolated for kurtosis scans (n=4, representative images are shown). The diffusivity and kurtosis were analyzed from mice treated with either PBS (control) or CL656 (test). Both parameters were assessed in the ipsilateral (ILS) and contralateral (CLS) striatum (n=4, * = $p<0.05$).

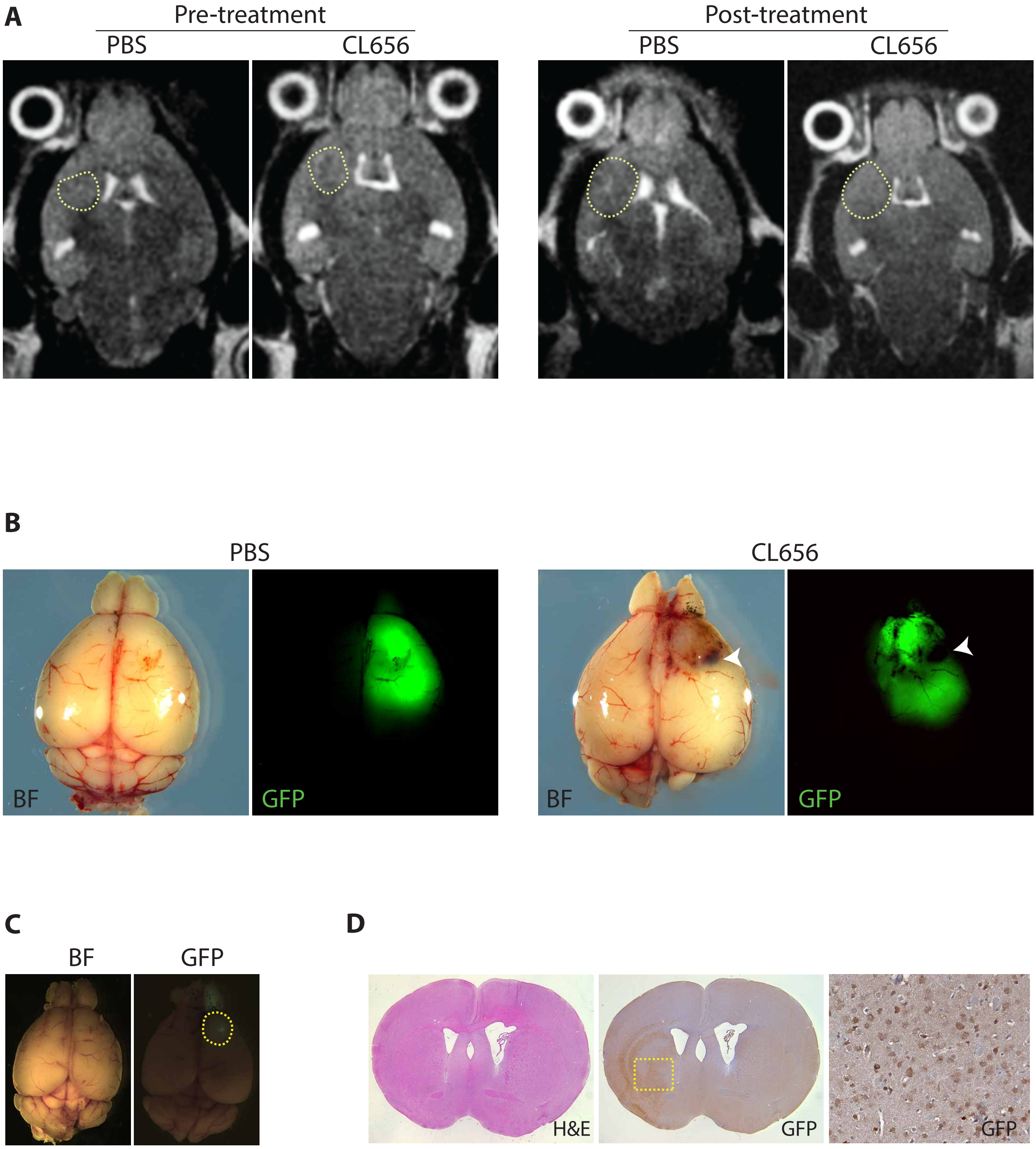
A
Pre-treatment
Post-treatment
PBS
CL656
PBS
CL656
B
PBS
CL656
BF
GFP
BF
GFP
C
BF
GFP
D
H&E
GFP
GFP

Fig S10

**Figure S10: Accelerated tumor progression by loss of Nf1**

A) GBM tumors with mutations in Nf1, Pten, and Trp53 were scanned 10 weeks after induction and post-treatment with CL656. A dashed yellow line marks the tumors (n=10).

B) Nf1 deficient tumors from PBS or CL656-treated mice were assessed by bright field (BF) and fluorescence microscopy (GFP). White arrowheads marks the hemorrhage (n=10, representative images are shown).

C) Mice were injected with AAV expressing GFP and 3 months post injection brains were harvest. Expression of GFP was assessed by fluorescence microscopy (GFP, marked by dash yellow line) (n=3, representative images are shown).

D) Brain sections were stained by H&E and GFP (brown stained). The dash box marks the area of magnification (n=3, representative images are shown).

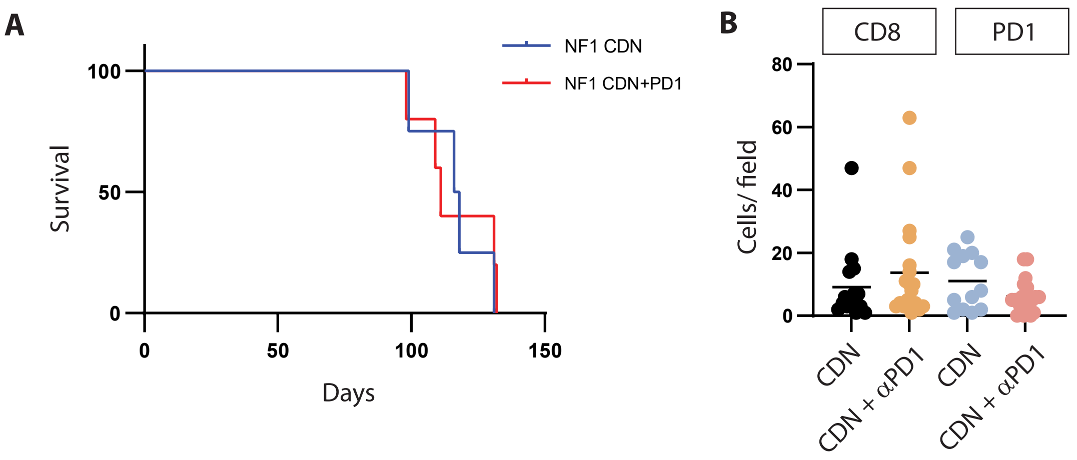
A
NF1 CDN
NF1 CDN+PD1
Survival
100
50
0
0
50
100
150
Days
B
CD8
PD1
Cells/ field
80
60
40
20
0
CDN
CDN + αPD1
CDN
CDN + αPD1

Fig S11

**Figure S11: Combination of CL656 and anti-PD1 treatment of Nf1 deficient mice**

Mice with Nf1 mutated GBM received for one-week CL656 treatment followed by further two-weeks treatment with either CL656 or CL656 and anti-PD1.

A) The survival of CL656- or CL656 and anti-PD1-treated mice was assessed at the humane endpoint (n=5).

B) Quantification of CD8 and PD1 positive cells in Nf1 deficient tumors from mice treated with CL656 or CL656 and anti-PD1 (n=15 fields).

| Primers used for qRT-PCR | | |
|---|---|---|
| **Genes** | | |
| **Isg15** | Taqman: Mm01705338 | |
| **Isg56** | accatgggagagaatgctgat | gccaggaggttgtgcatc |
| **Cxcl10** | cgatgacgggccagtgagaatg | tcaacacgtgggcaggataggct |
| **Gapdh** | CAA TGT GTC CGT CGT GGA | GAT GCC TGC TTC ACC ACC |
| **Vegfa** | Mm00437306 | |
| **Eno1** | Mm01619597 | |
| **Brip3** | Mm00833810 | |
| **Actin** | Mm02619580 | |

| Primers used for PCR | | |
|---|---|---|
| **Genes** | | |
| **Mouse** | **Forward primer** | **Reverse primer** |
| Pten | CTCCCTGGAGTGAAGAGCAC | GTGTGCCTAGCACCTACTCC |
| Rb1 | TAGGGCCTGGGTTGCTTCTA | TTGGCAACTGCTCACACACT |
| Tp53 | ATAGAGACGCTGAGTCCGGT | CAAAGAGCGTTGGGCATGTG |
| Nf1 | ACATGCAAGTGGCTGGATCA | TGGAACAAATCTTTGGGAAAAGAGG |

| Protospacers used for Cas9 targets | |
|---|---|
| **Mouse** | **guide RNA (protospacer)** |
| Pten | GCTTTACAGTGAATTGCTGC |
| Rb1 | AGCATTATCAACCTTGGTAC |
| Tp53 | GTGTAATAGCTCCTGCATGG |
| Nf1 | GCTGCAGCCAAGAGCTCTTG |
